# Comparing Brain-Behavior Relationships Across Dimensional, Tail-Sampled, and Propensity-Matched Models

**DOI:** 10.1101/2025.05.23.655740

**Authors:** K. Murtha, L. Dorfschmidt, R.A.I. Bethlehem, K. Kang, S. Vandekar, A.F. Alexander-Bloch, R. Waller, J. Seidlitz

## Abstract

Large population cohorts are needed to perform brain-wide association studies (BWAS), with evidence that sampling from the tails of a distribution increases effect sizes and improves reproducibility. However, studies rarely compare how variability in sample sociodemographic characteristics relates to imaging or behavioral phenotypes within BWAS. To address this gap, we derived estimates for brain-behavior associations using multivariate regression models, comparing effect sizes for dimensional, tail-sampled, and propensity matched groups. Data were obtained from the Adolescent Brain Cognitive Development (ABCD) Study®. The independent variables were brain structural imaging phenotypes, with a range of biological and psychological outcomes as dependent variables. We found expected increases in the magnitude of effect sizes moving from full-sample dimensional models to tail-sampled group-based models. However, findings for the propensity-matched group models suggested a non-uniform impact on BWAS (i.e., both increased and decreased effect sizes). Results suggest that sampling from the tails of the distribution of measures of brain structure generally increases effect sizes across biological, clinical, and cognitive outcomes.

## Main

Much research has been devoted to uncovering the neurobiological underpinnings of different mental health disorders, including through the use of non-invasive magnetic resonance imaging (MRI; Buckholtz & Meyer-Lindenberg, 2012; Chen et al., 2022; Durham et al., 2023; Xia et al., 2018). Brain-wide association studies (BWAS) are an important tool to describe the relationship between MRI-derived measures of brain structure and function and various clinical, psychological, and cognitive phenotypes, as well as environmental risk exposures (Marek et al., 2022; Owens et al., 2021). Recently, however, concerns have been raised about the reproducibility and replicability of BWAS studies. These concerns have centered on the need for larger sample sizes (Marek et al., 2022), as well as emphasizing the low prediction accuracy (Schulz et al., 2024) and small effect sizes (Reddan et al., 2017; Szucs & Ioannidis, 2020) of extant literature. Various methodological improvements have been proposed to address the reproducibility and replicability of BWAS studies conducted in both small and large samples (He et al., 2022; Rosenberg & Finn, 2022), including increasing the standard deviation of an independent variable by sampling from the tails of its distribution (Kang et al., 2024).

However, prior work has yet to consider how individual differences in sociodemographic characteristics within and between samples might impact reported brain-behavior associations. For example, several prior studies have linked the development of cognitive skills to the maturation of the executive system across childhood and adolescence (Keller et al., 2023; Luna, 2009; Satterthwaite et al., 2013; Shamosh et al., 2008). Concurrently, a large literature demonstrates that socioeconomic disadvantage measured at the individual, family, and neighborhood levels is related to both cognitive development (Hackman et al., 2015; Last et al., 2018; Lawson & Farah, 2017; Noble et al., 2007) and the maturation of brain executive function systems (Farah, 2017; Mackey et al., 2015; Noble et al., 2015). This pattern of relationships among risk factors and outcomes creates a “third variable problem,” (Westfall & Yarkoni, 2016) in which the effect of a third variable (i.e., socioeconomic status) can obscure the relationship between hypothesized exposures (i.e., maturation of the executive function network in the brain) and outcome variables (i.e., cognitive functioning), which has rarely been considered in BWAS. The third variable problem differs from collider bias, wherein conditioning variables on a common effect induces an artificial association between them (Holmberg & Andersen, 2022). This third variable problem is also evident in many studies that have examined the relationships between brain structure or function and symptoms of psychopathology (Gur et al., 2019; Qiu & Liu, 2023) or biological outcomes, such as birth weight or gestational age at birth (El Marroun et al., 2020; Finch, 2003).

To address the third variable problem, studies commonly statistically control for sociodemographic characteristics that might confound the relationship between predictor and outcome variables within BWAS that employ regression analyses. Such covariates typically include age, sex, socioeconomic status of individuals, or motion measures evaluated during the MRI scanning. However, it is possible that controlling for covariates that are not true confounders can bias the estimates obtained between brain and behavior (Wysocki et al., 2022). Likewise, typical de-confounding tools used in machine learning models, such as regressing out the effects of age and sex, and using cleaned variables, may not adequately remove the signal of a given covariate as they cannot account for non-linear effects (Benkarim et al., 2022).

The most rigorous statistical approach to address causality is through experimental designs, such as randomized controlled trials (RCT), which are not feasible in studies examining brain-behavior relationships. To approximate causal inference, propensity matching methods can be employed to mimic the conditions of a RCT by reducing bias when comparing naturally occurring groups in observational research (D’Agostino, 1998). Propensity scores are estimated by calculating the likelihood of being in an experimental versus control condition based on a set of observed covariates using logistic regression. Those scores are used in a variety of matching techniques to select groups of control participants who “match” individuals in the experimental condition across covariates included in matching procedures (Austin, 2011, 2014). Ultimately, this approach can facilitate comparisons between groups who are identical on sociodemographic characteristics that might be otherwise related to the independent or dependent variables of study, thus approximating the bias reduction that would be made possible through random assignment (Rosenbaum & Rubin, 1983). When applied to BWAS, propensity matching based on the tails of a distribution of MRI-derived brain phenotypes may allow for more parsimonious control of unwanted confounders by selecting participants who have similar sociodemographic profiles that could otherwise differ between groups. While propensity score matching is typically applied to the independent variable of interest, the purpose of this work was to better elucidate the differential contributions of sociodemographic variables in the context of BWAS, and thus has more implications for reporting associations rather than study design.

Using data from the Adolescent Brain Cognitive Development (ABCD) Study®, which is one of several large imaging cohorts used frequently to examine BWAS relationships, our goal was to test whether using effect sizes obtained for several brain-behavior associations varied across three different methods: full sample (i.e., dimensional), tail-sampled groups, and propensity-matched groups. We focused on morphological brain measures based on their interpretability and extensively documented relationships (Casey et al., 2000; Gale-Grant et al., 2021; Mewton et al., 2022) with various clinical, cognitive, and biological outcomes. We assessed the following morphological brain measures as independent variables: cortical gray matter volume (cGMV) – including specific left- and right-hemisphere regional volumes, subcortical gray matter volume (sGMV), white matter volume (WMV), and ventricular cerebrospinal fluid volume (CSF). Our independent variables featured a host of biological and psychological outcomes, including birthweight and gestational age, symptoms of psychopathology, and performance on cognitive tests. First, we tested dimensional associations in the full study sample, conventionally controlling for covariates. Second, we repeated analyses using groups that were sampled from the tails of the morphological brain measure distributions, selecting participants with scores falling one median-absolute deviation above or below the sample median, controlling for covariates. Finally, we created propensity matched samples, selecting participants with high and low representations of each morphological brain measure from the median-absolute deviation samples who were matched on age, sex, pubertal stage, race, ethnicity, parental education, household income, and image quality during structural imaging as measured by Euler number. To ensure comparability across model types, we use the Robust Effect Size Index (RESI), which is less sensitive to outliers and allows application to skewed or non-normally distributed data (compared to traditional effect size measures such as Cohen’s *d*) (Vandekar et al., 2020). Across analyses, we hypothesized that effect sizes would increase in tail-sampled models, and either increase or decrease in propensity matched models, given the minimal variation in sociodemographic profiles of included participants that could otherwise have been related to dependent and independent variables. Establishing this reporting framework represents an important foundation for future studies’ interpretations of findings in BWAS.

## Results

### Sample Characteristics and Demographics

Using baseline structural MRI, behavioral, and parent report data from the 5^th^ release of the ongoing Adolescent Brain Cognitive Development (ABCD) study, we created complete, tail-sampled, and propensity matched samples to model relationships between four brain tissue types and biological, clinical, and cognitive outcomes. Descriptive statistics for each sampling strategy for each tissue type, as well as any group differences on matching variables are presented in **Tables S1-S4**. Groups matched on different morphological brain features did not differ significantly from each other on most matching variables, while matched groups differed significantly from tail-sampled groups on a variety of measures, including household income and parental education (**Appendix 1**). Propensity matching procedures yielded adequate balance, where standardized mean differences of covariates between groups were all <.10 across all tissue types (**Figure S1**).

### Comparing findings across methods for brain-biological outcomes

RESIs in models predicting biological outcomes across tissue types most frequently showed increases from dimensional to tail-sampled models, which were not significantly reduced when we used propensity matching methods (**Table 1**, **Figure 1**). Higher whole-brain cGMV was consistently associated with older gestational age across dimensional, tail-sampled, and propensity matched models. Higher cGMV was also associated with greater birth weight, with increasing effect sizes from dimensional to tail-sampled models, but not propensity matched models (**Table S5**). Likewise, both higher sGMV and higher WMV were associated with higher gestational age and higher birth weights, with increasing effect sizes between dimensional and tail sampled models, but not propensity-matched models. However, for BMI, we found that sGMV and WMV volumes were associated with higher BMI only in tail-sampled and propensity matched models, but not in dimensional models (**Tables S6, S7**). Higher CSF was associated with higher birth weight across all modeling strategies (**Table S8**). To establish the robustness of this pattern of findings, we showed that effect sizes were relatively stable across differently sized randomly selected samples (**Figure S2).**

**Figure 1.**
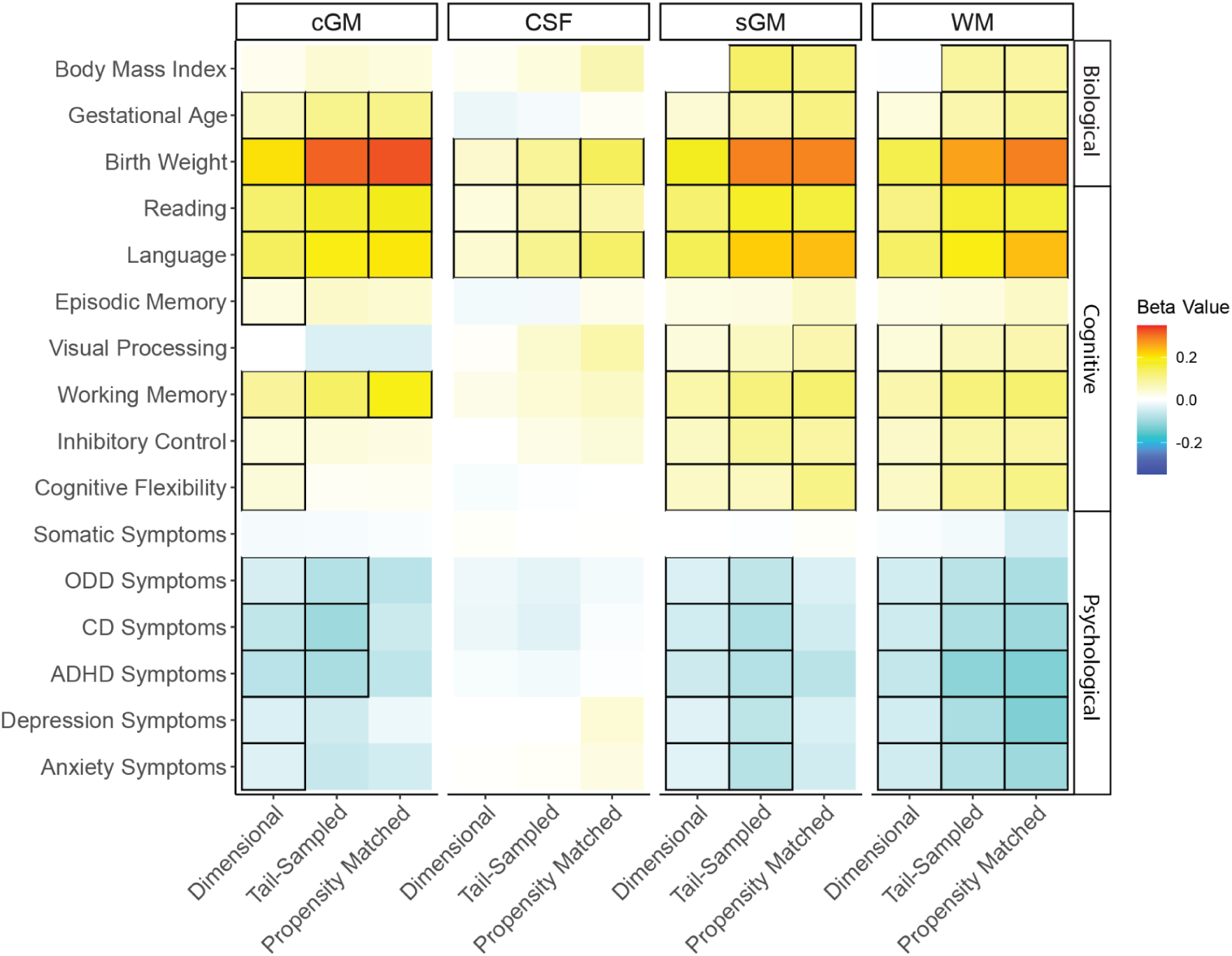
Heatmap figure of RESI values for biological, cognitive, and psychological outcomes across cGMV, sGMV, WMV, and CSF whole brain tissue types. Significant associations are outlined in black.

**Table 1.** Regression results for models predicting biological outcomes using cGMV, WMV, sGMV, and CSF across 3 modeling frameworks, controlling for covariates.

|  | cGMV |  | WMV |  | sGMV |  | CSF |  |
| --- | --- | --- | --- | --- | --- | --- | --- | --- |
|  | B (SE) | RESI | B (SE) | RESI | B (SE) | RESI | B (SE) | RESI |
| **Dimensional Models** |  |  |  |  |  |  |  |  |
| BMI | 0.94, (0.64) | 0.019 | -0.31, (0.99) | -0.004 | 0.11, (1.3) | 0.001 | 3.45, (2.92) | 0.015 |
| Gestational Age | 0.23, (0.05) | 0.064*** | 0.12, (0.04) | 0.033* | 0.15, (0.05) | 0.041** | -0.07, (0.04) | -0.021 |
| Birthweight | 1.05, (0.07) | 0.205*** | 0.76, (0.07) | 0.152*** | 0.88, (0.07) | 0.173*** | 0.24, (0.07) | 0.047*** |
| **Tail Sampled Models** |  |  |  |  |  |  |  |  |
| BMI | 0.37, (0.21) | 0.042 | 0.73, (0.2) | 0.088*** | 1.17, (0.21) | 0.134*** | 4.53, (3.65) | 0.031 |
| Gestational Age | 0.23, (0.05) | 0.105*** | 0.16, (0.05) | 0.076* | 0.18, (0.05) | 0.086** | -0.03, (0.06) | -0.012 |
| Birthweight | 1, (0.08) | 0.314*** | 0.79, (0.08) | 0.262*** | 0.86, (0.08) | 0.289*** | 0.31, (0.08) | 0.097** |
| **Propensity Matched Models** |  |  |  |  |  |  |  |  |
| BMI | 0.27, (0.28) | 0.033 | 0.68, (0.24) | 0.087* | 1.04, (0.28) | 0.119*** | 0.59, (0.31) | 0.073 |
| Gestational Age | 0.21, (0.07) | 0.11* | 0.2, (0.06) | 0.099** | 0.23, (0.06) | 0.114** | 0.02, (0.08) | 0.012 |
| Birthweight | 0.91, (0.1) | 0.324*** | 0.84, (0.09) | 0.292*** | 0.81, (0.09) | 0.287*** | 0.46, (0.13) | 0.145** |

Results were largely similar when we examined specific brain regions (**Table S9**, **Figure 2**). Across the left and right hemispheres, no regions significantly predicted BMI in dimensional models (all p_FDR_ > 0.05). Some regions saw significant associations with BMI in tail-sampled models, however most of these individual relationships were reduced when modeled in propensity matched samples. Regional measures of cGMV were consistently related to higher gestational age and were strongest in dimensional and propensity matched models. Finally, regional measures of cGMV were related to higher birth weights, with higher RESIs in propensity matched models (**Figure S3**).

**Figure 2.**
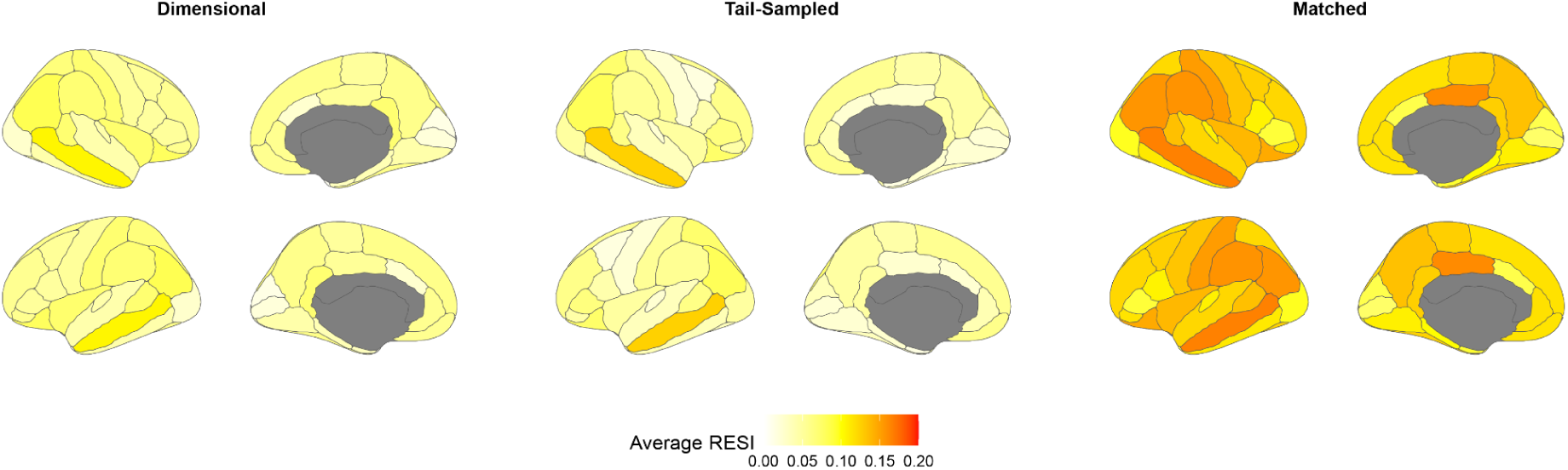
Cortical brain maps showing associations of regional measures of cGMV with biological outcomes across dimensional, tail sampled and propensity matched models. The average of the absolute value of each RESI was taken across 3 biological outcomes.

### Comparing findings across methods for brain-cognitive outcomes

Patterns of RESIs in models predicting cognitive outcomes differed across tissue type, but overall showed positive relationships between tissue volume and performance across cognitive domains. Across all modeling strategies, greater cGMV volume was related to better performance across all cognitive measures, except for episodic memory and visual processing (**Table 2**, **Figure 1**). For reading, language, and working memory, greater cGMV was related to higher scores across dimensional, tail-sampled, and propensity matched methods (**Table S10**). For inhibitory control and cognitive flexibility, greater cGMV was related to higher scores in dimensional models, but not in tail-sampled or propensity matched models. Effect sizes were largely consistent across different size randomly selected samples (**Figure S2).**

**Table 2.**
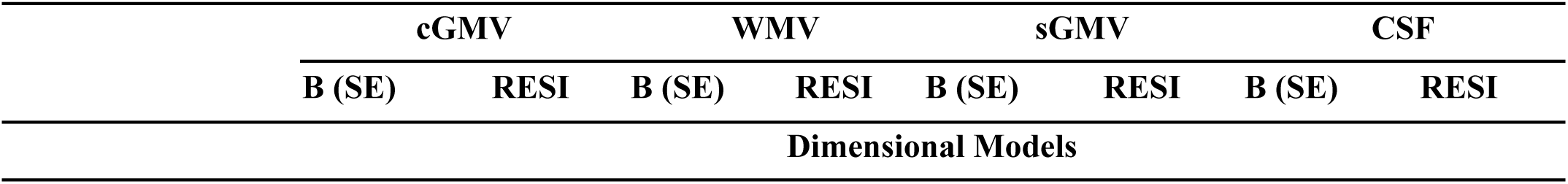
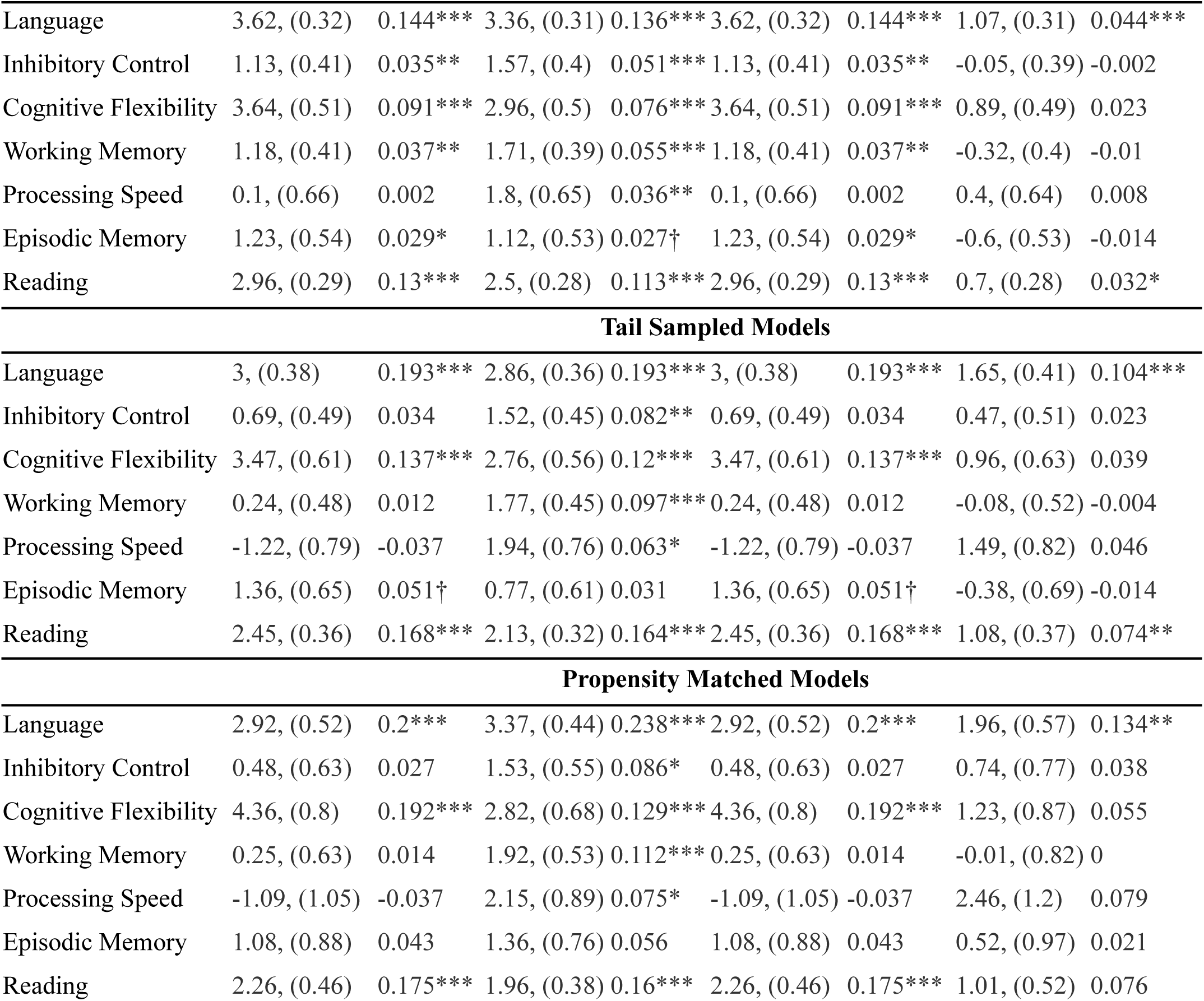
Regression results for models predicting cognitive outcomes using cGMV, WMV, sGMV, and CSF across 3 modeling frameworks, controlling for covariates.

Greater sGMV volume was related to higher reading, language, working memory, and inhibitory control scores across all modeling strategies (Table S11). For visual processing, higher scores were only related to greater sGMV in dimensional models, but not tail-sampled or propensity matched models. For cognitive flexibility, higher scores were related to greater sGMV in dimensional and propensity matched models, but not tail-sampled models.

Higher WMV was related to higher scores on reading, language, working memory, inhibitory control, and cognitive flexibility across all modeling strategies (Table S12). Visual processing was related to WMV in dimensional, and tail-sampled models, but not propensity matched models. Episodic memory was not related to WMV across any modeling strategies. Higher CSF was related to language across all modeling strategies. Higher CSF was related to reading in dimensional and tailed sampled models, but not in propensity matched models (Table S13). CSF was not significantly related to scores on any other cognitive dimensions.

Relationships between regional measures of cGMV and cognitive variables were largely consistent with patterns of results from whole-brain measures (**Figure 3**, Table S14). Across the left and right hemispheres, regional measures of cGMV were consistently related to scores on reading, language, and working memory, and were strongest in propensity matched models. In tail sampled models, regional measures of cGMV were inconsistent, with some regions showing negative relationships (Figure S4). Many of the regional relationships that were significantly related to domains including cognitive flexibility, inhibitory control, visual processing, or episodic memory in either dimensional or tail-sampled models were rendered non-significant in propensity matched models.

**Figure 3.**
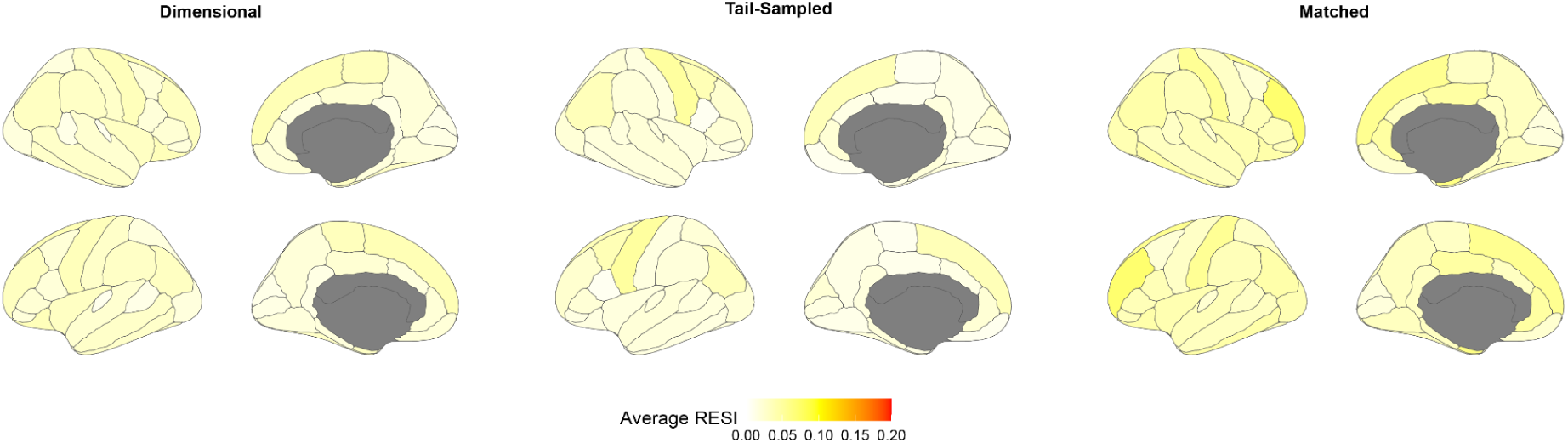
Cortical brain maps showing associations of regional measures of cGMV with cognitive outcomes across dimensional, tail sampled and propensity matched models. The average of the absolute value of each RESI was taken across 7 cognitive outcomes.

### Patterns of results for symptoms of psychopathology

Patterns of RESIs in models predicting psychological outcomes differed across tissue types, but overall showed that lower tissue volume related to higher symptom counts (**Table 3**, **Figure 1**). Lower cGMV was significantly associated with more symptoms of Oppositional Defiant Disorder (ODD), Conduct Disorder (CD), and Attention Deficit Hyperactivity Disorder (ADHD) in dimensional and tail-sampled models (**Table S15**). Lower cGMV was associated with more symptoms of anxiety and depression in dimensional models, but not in either tail sampled or propensity matched models. cGMV was not significantly associated with somatic symptoms across any modeling strategy. Effect sizes were largely consistent across different size randomly selected samples (**Figure S2).**

**Table 3.** Regression results for models predicting psychological outcomes using cGMV, WMV, sGMV, and CSF across 3 modeling frameworks, controlling for covariates.

|  | cGMV |  | WMV |  | sGMV |  | CSF |  |
| --- | --- | --- | --- | --- | --- | --- | --- | --- |
|  | B (SE) | RESI | B (SE) | RESI | B (SE) | RESI | B (SE) | RESI |
| **Dimensional Models** |  |  |  |  |  |  |  |  |
| Somatic Symptoms | -0.07, (0.07) | -0.013 | -0.04, (0.07) | -0.007 | 0, (0.07) | 0 | 0.04, (0.07) | 0.007 |
| ODD Symptoms | -0.32, (0.09) | -0.044** | -0.34, (0.09) | -0.048*** | -0.28, (0.09) | -0.038** | -0.14, (0.09) | -0.02 |
| CD Symptoms | -0.53, (0.1) | -0.066*** | -0.4, (0.1) | -0.052*** | -0.38, (0.1) | -0.048*** | -0.17, (0.1) | -0.022 |
| ADHD Symptoms | -0.77, (0.13) | -0.073*** | -0.66, (0.13) | -0.064*** | -0.57, (0.13) | -0.054*** | -0.1, (0.13) | -0.01 |
| Depression Symptoms | -0.26, (0.09) | -0.037** | -0.33, (0.09) | -0.048*** | -0.25, (0.09) | -0.034* | 0.01, (0.09) | 0.002 |
| Anxiety Symptoms | -0.32, (0.11) | -0.036* | -0.41, (0.11) | -0.049*** | -0.3, (0.11) | -0.033* | 0.05, (0.11) | 0.006 |
| **Tail Sampled Models** |  |  |  |  |  |  |  |  |
| Somatic Symptoms | -0.04, (0.08) | -0.012 | -0.05, (0.08) | -0.015 | -0.02, (0.08) | -0.005 | -0.01, (0.09) | -0.001 |
| ODD Symptoms | -0.35, (0.11) | -0.079** | -0.32, (0.11) | -0.074** | -0.3, (0.11) | -0.068* | -0.14, (0.12) | -0.03 |
| CD Symptoms | -0.48, (0.12) | -0.101*** | -0.38, (0.11) | -0.084** | -0.38, (0.12) | -0.082** | -0.2, (0.15) | -0.034 |
| ADHD Symptoms | -0.57, (0.15) | -0.09*** | -0.74, (0.15) | -0.116*** | -0.51, (0.15) | -0.08** | -0.11, (0.17) | -0.016 |
| Depression Symptoms | -0.21, (0.1) | -0.051 | -0.37, (0.1) | -0.087** | -0.31, (0.11) | -0.07* | -0.01, (0.12) | -0.002 |

| Anxiety Symptoms | -0.31, (0.12) | -0.06† | -0.41, (0.13) | -0.08** | -0.4, (0.13) | -0.076* | 0.06, (0.15) | 0.01 |
| --- | --- | --- | --- | --- | --- | --- | --- | --- |
| Propensity Matched Models |  |  |  |  |  |  |  |  |
| Somatic Symptoms | -0.03, (0.11) | -0.009 | -0.14, (0.1) | -0.045 | 0.02, (0.1) | 0.007 | 0.01, (0.13) | 0.004 |
| ODD Symptoms | -0.3, (0.14) | -0.073 | -0.36, (0.13) | -0.088† | -0.16, (0.13) | -0.038 | -0.06, (0.18) | -0.014 |
| CD Symptoms | -0.25, (0.16) | -0.057 | -0.41, (0.13) | -0.099** | -0.21, (0.13) | -0.05 | -0.04, (0.23) | -0.007 |
| ADHD Symptoms | -0.41, (0.21) | -0.068† | -0.76, (0.19) | -0.126*** | -0.46, (0.19) | -0.074† | -0.03, (0.26) | -0.005 |
| Depression Symptoms | -0.08, (0.14) | -0.02 | -0.54, (0.13) | -0.128*** | -0.19, (0.14) | -0.042 | 0.19, (0.19) | 0.039 |
| Anxiety Symptoms | -0.23, (0.17) | -0.048 | -0.47, (0.14) | -0.1* | -0.26, (0.16) | -0.049 | 0.16, (0.22) | 0.028 |

Lower sGMV was associated with more symptoms of ODD, CD, ADHD, depression, and anxiety with modest increases in effect sizes moving from dimensional to tail sampled models. However, sGMV was unrelated to symptoms of psychopathology in propensity matched models (Table S16). Lower WMV was associated with all symptom measures except for somatic symptoms across all modeling strategies, with slight increases in effect sizes, (Table S17). CSF was not significantly associated with any symptom measures (Table S18).

Relationships between regional measures of cGMV and cognitive variables were consistent with patterns of results from whole-brain measures (**Figure 4**, Table S19). Across the left and right hemispheres, lower regional cGMV measures were consistently related to more symptoms of ADHD, CD, ODD, depression, and anxiety, with small effect sizes (i.e., likely contributing to whole brain cGMV effects). The number of regions that were significantly related to symptoms of psychopathology in tail-sampled models decreased, while effect sizes for those that remained were larger, specifically for ADHD, CD, and ODD symptoms (Figure S5). While very few regions were significantly related to symptoms in propensity matched models, those that did evidenced stronger effect sizes.

**Figure 4.**
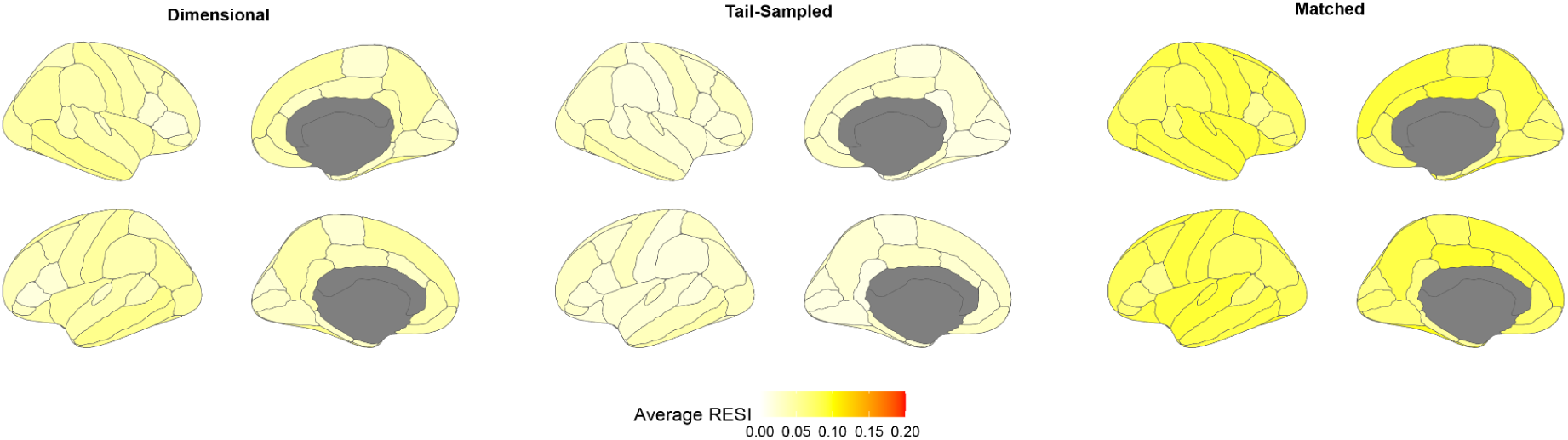
Cortical brain maps showing associations of regional measures of cGMV with psychological outcomes across dimensional, tail sampled and propensity matched models. The average of the absolute value of each RESI was taken across 6 psychological outcomes.

## Discussion

We examined how study design relates to findings for brain-behavior associations, seeking to extend prior work focused on increasing the replicability of BWAS through use of different sampling methods (Kang et al., 2024; Marek et al., 2022). Overall, our results suggest increasing effect sizes when modeling the relationship between whole-brain measures of cGMV, sGMV, WMV, and CSF and various biological, cognitive, and psychological outcomes. However, this pattern of findings was not universally consistent, particularly when applying more stringent propensity matching procedures to compare groups. We discuss the implications of these findings for future BWAS.

Overall, our findings echo many reported BWAS findings from the ABCD study and other large neuroimaging samples. We found that greater gray and white matter volume was consistently related to better performance across cognitive domains (Magistro et al., 2015; Michel et al., 2024; Muetzel et al., 2015; Tamnes et al., 2010). Likewise, lower gray and white matter volume was largely related to more symptoms across internalizing and externalizing dimensions of psychopathology (Durham et al., 2021; Kaczkurkin et al., 2019; Li et al., 2020; Neumann et al., 2020; Snyder et al., 2017). We also replicate prior findings linking biological measures at birth to brain development, with higher gray and white matter volume both related to higher birth weight (de Kievet et al., 2012; Walhovd et al., 2024) and older gestational age at birth (Ma et al., 2022; Nath et al., 2023).

However, while many of the model comparisons evidenced expected increases in the magnitude of effect sizes between full-sample dimensional and tail-sampled models (Kang et al., 2024), there were important differences moving to the propensity-matched models, where more stringent adjustment for sociodemographic and methodological variables had a non-uniform impact on BWAS across tissue types. By reducing the variability of covariates and making them more independent to measures of brain structure, propensity score matching increased or decreased effect sizes depending on their relationship with the brain variable and outcome variable. For the cognitive measures, relationships for whole brain gray and white matter volume were largely consistent moving from tail-sampled to propensity matched models. This pattern of results demonstrates the criticality of brain structure differences for understanding cognitive functioning, over and above the influence sociodemographic characteristics that have been consistently linked both to brain structure (Brito & Noble, 2014; Farah, 2017) and cognition (Duncan & Magnuson, 2012; Noble et al., 2007). Exceptions to this pattern of findings from the cognitive domain were for the associations between cGMV and inhibitory control and cognitive flexibility, with significant effect sizes in dimensional but not tail-sampled or propensity-matched group models. These exceptions suggest that specific cognitive functions are associated with specific morphological profiles and may have unique associations with sociodemographic variables.

Likewise, when it came to examining psychopathology, we documented key differences between dimensional versus tail-sampled and propensity matched models, such that symptoms of both internalizing and externalizing psychopathology were only related to cGMV or sGMV in dimensional and tail-sampled models. A particularly striking example of this trend was for symptoms of ADHD, ODD, and CD, where effect sizes decreased in propensity matched models, suggesting that relationships between morphological brain measures and externalizing disorders may be explained by shared etiological risk factors captured by measures of socioeconomic status or parental education (Peverill et al., 2021). These findings suggest that prior BWAS studies that have emphasized links between brain structure and psychopathology without sensitivity analyses or more stringent control for sociodemographic characteristics, may have overstated the evidence for links between psychopathology and brain structure (Farah, 2017; Gur et al., 2019; Peverill et al., 2021). That is, our pattern of findings emphasizes that much of the variance captured in the dimensional models linking brain structure to psychopathology appears to be related to sociodemographic characteristics, which has important implications for translation and policy (Perkins et al., 2024).

Together, our results highlight the need for greater attention to the meaningfulness of effect sizes and use of more stringent methods to account for sociodemographic characteristics within BWAS. The current work extends the discussion of new methods for more robust statistical reporting, including robust effect size indices (Kang et al., 2024) by considering the ways in which relationships in BWAS studies may be affected by shared etiological risk factors. By conducting sensitivity analyses that parametrically varied sample size, we ruled out cohort-size related effects, which allows for a more direct evaluation of effect sizes across propensity matched and tail-sampled models. Further, leveraging methods like brain charting allow us to harmonize morphological brain phenotypes to ensure a more rigorous quantification of individual differences in brain structure (Bethlehem et al., 2022). In concert, these approaches will be useful in increasing robustness of BWAS studies and may ultimately contribute to the design and implementation of precision medicine solutions.

Second, simply statistically adjusting for factors (e.g., adjusting for covariates in analyses) that differ between groups using traditional modeling methods violates key statistical assumptions, including the independence of covariates and group assignment (Miller & Chapman, 2001) and can distort relationships between collinear covariates, resulting in inaccurate or unreproducible results (Hoyle et al., 2023; Swilley-Martinez et al., 2023; Wysocki et al., 2022). For example, consistent with prior work, children from the “high” end of the tails of the various morphological measures were from households with significantly higher incomes and had parents reporting higher educational attainment (King et al., 2019; Noble et al., 2012; Taylor et al., 2020). In contrast, at the “low” end of the tail were from households with lower income and had parents who reported lower educational attainment, consistent with studies showing that individuals from more advantaged backgrounds show higher brain volume across tissue types (Rakesh et al., 2022). To examine BWAS across such groups would clearly violate statistical assumptions. By creating propensity-matched groups, we demonstrate that it is possible to avoid findings that reflect potentially spurious group differences, particularly if propensity matching is used as an additional step to establish the robustness of findings. However, propensity matching is not without limitations, including the potential inexact or incomplete matching, which may result in residual confounding or a lack of generalizability or results (Sainani, 2012), with Mahalanobis Distance Matching or Coarsened Exact Matching representing two additional methods to address the third variable problem we describe (King & Nielsen, 2019).

Strengths of the current study include the range of morphological, biological, psychological, and cognitive variables, as well as replication across whole brain and regional measures. However, findings should be considered alongside key limitations. First, while propensity matching allowed us to rigorously account for sociodemographic and methodological factors, children in the low propensity matched group often differed significantly from individuals in low tail-sampled groups on measures of household income and parental education. That is, by sampling from the tails of the distribution for a given structural phenotype, and because these phenotypes are related to many variables included in our matching procedures (e.g., parental education, socioeconomic status), our approach may have artificially constructed groups that were not fully representative of the whole cohort. Thus, while the associations reported here between measures of brain volume and biological, psychological, and cognitive variables cannot be attributed to differences in characteristics between propensity matched groups, individuals in our matched groups may not be representative of the general population. Second, sample size differed significantly across dimensional, tail-sampled, and propensity matched groups, which could have contributed to differences in statistical significance across modeling strategies (Sullivan & Feinn, 2012). However, effect sizes were relatively stable when repeatedly tested across random samples of various sizes. Third, while we examined a variety of different measures of brain structure, future work could apply our methods to evaluate relationships between brain function (i.e., task-based cerebral blood flow, connectivity, topography) or multi-ancestry polygenic indices of genetic risk for biological, cognitive, or clinical outcomes.

We established the utility of a propensity matching approach as a rigorous reporting framework for BWAS. As changes in sampling and modeling strategies can have varied effects on BWAS across a range of outcomes, studies should include a range of different approaches to test the robustness of findings within any analytic pipeline. To facilitate reproducible reporting of BWAS using propensity score matching in the widely-used ABCD study, our tail-sampled and propensity-matched groups will be made available through future releases of the ABCD data and our code to reproduce the groups for investigators is already available at https://github.com/krmurtha1/BWAS_matching

## Methods

### Data Availability

We used structural imaging, behavioral, and parent report data from the fifth release of the ongoing Adolescent Brain Cognitive Development (ABCD) study, recruitment and sampling strategies for which have been described in detail previously (Garavan et al., 2018).

### Measures

#### Matching Variables

Parents completed the Parent Longitudinal Demographic Questionnaire at the baseline assessment, which included questions about child age, sex, race, ethnicity, parental education, and household income (Barch et al., 2018; Hamilton et al., 2011). Pubertal status was assessed with the Youth Pubertal Development Scale (Barch et al., 2018; Petersen et al., 1988). MRI Motion was assessed using Euler number, a measure of image quality during T1 and T2 scans calculated by Freesurfer (Rosen et al., 2018; Tobe et al., 2024).

#### Morphological Brain Measures

Details of data processing for the imaging-derived phenotypes for the ABCD study, as well as the population normative models (“brain charts”), are described in previous work (Bethlehem et al., 2022). Briefly, we used whole-brain centile scores for 4 main tissue types, including total cortical gray matter volume (cGMV), subcortical gray matter volume (sGMV), white matter volume (WMV), and cerebrospinal fluid volume (CSF). Using the centile scores, compared to the raw volumes, allows for robust harmonization across technical variables (i.e., site) and demographic variables (age and sex) and for comparison in statistical effects across phenotypes. Total cGMV, sGMV, WMV and CSF were estimated from T1- and T2-weighted scans using FreeSurfer version 6.1. Centile scores were estimated relative to reference curves of developmental milestones, to arrive at individual scores (between 0-1) that can be interpreted similarly to a traditional growth chart. Exploratory analyses were performed using centile scores calculated at the regional level, which were mapped across the 68 cortical regions (34 per hemisphere) of the Desikan-Killiany atlas (Desikan et al., 2006).

#### Biological Outcomes

Gestational age and birthweight were obtained from the Developmental History Questionnaire. Gestational age was calculated using parent report on how many weeks premature the child was born. Following recommendations from prior work on this variable in ABCD, gestational age was coded into 5 groups, corresponding to adolescents born at less than 33 weeks, 34–35 weeks, 36 weeks, 37–39 weeks, and 40 weeks of gestation (Ma et al., 2022). Birthweight was reported in pounds and ounces. BMI was calculated using participants height and weight.

#### Cognitive Outcomes

The ABCD study includes a variety of cognitive tasks from the NIH Cognition Battery Toolbox, which index language and reading ability, episodic and working memory, as well as dimensions of executive function, including inhibitory control, cognitive flexibility, and processing speed (Luciana et al., 2018; Thompson et al., 2019). Age-uncorrected task scores were used in the current analyses.

#### Child Psychopathology

To measure broad domains of child psychopathology, we used the DSM-oriented subscales of the Child Behavior Checklist from baseline, focusing on ADHD, ODD, CD, Anxiety, Depression, and Somatic Symptoms. Items are scored on a 3-point scale (0=absent, 1=sometimes, 2=occurs often).

### Sample Identification

First, dimensional, median absolute deviation, and propensity matched samples were identified **(Figure S6**). We used complete case analysis (CCA) to address missing data, and participants with missing data on covariates were excluded, following recommendations against matching on estimated or imputed data (Granger et al., 2019; Knol et al., 2010). Participants with complete versus incomplete differed significantly on measures of cGMV, sGMV, and WMV, but effect sizes were negligible (**Figure S4**). One sibling at random was excluded from the sample to account for family level effects. For each tissue type, we identified subjects who scored one median-absolute deviation above or below the sample median.

To create propensity matched samples of children with high versus low representation of each brain structure metric, we used the R Package “MatchIt” (Ho et al., 2011). Our matching features were age, sex, data collection site, parental education, combined household income, race, ethnicity, pubertal stage, and Euler number. We also included an age-by-sex interaction, to account for sex differences in development (Koolschijn & Crone, 2013). Participants were matched across groups using one-to-one nearest neighbor matching with Caliper distance=.10, with propensity scores estimated using logistic regression. In order to avoid a priori hypotheses about the direction of relationships between brain structure and outcomes of interest, we repeated matching procedures twice, first identifying individuals with high representations of each tissue type as the “treatment” group, and matching to a pool of “controls” with low representations, then identifying individuals with low representations as the “treatment” group and matching to a pool of “controls” with high representations. The final matched samples included participants who were identified when matching in both directions.

### Whole Brain Modeling Approach

To examine associations between morphological brain measures and biological, clinical and cognitive outcomes, we leveraged multivariate regression modeling using the Lavaan package in R (Rosseel, 2012), specifying correlated dependent variable models. First, in dimensional models, we tested the main effects of each tissue type on biological (BMI, Birth Weight, Gestational Age), clinical (all DSM-oriented subscales of the CBCL) and cognitive outcomes (all tasks from the NIH cognition toolbox) in 3 models (i.e., specifying each of these outcomes to be correlated). Second, in Median Absolute Deviation Models, we conducted the same analyses using binary group predictors (i.e.: high and low representations). Finally, we repeated these analyses in Propensity Matched samples. Multiple comparisons were corrected for across tissue types using FDR correction. All sociodemographic and methodological variables included in the propensity matching procedures were included as covariates across all models.

### Regional Modeling Approach

To better understand how the patterns of results with regard to cGMV were informed by regional differences, exploratory analyses were conducted by repeating our modeling approach using centile scores calculated at the regional level, which were mapped across the 68 cortical regions (34 per hemisphere). Each regional cGMV value was used as an independent variable across correlated dependent variable models as described above, tested within dimensional, tail-sampled, and propensity matched samples that were identified using whole-brain cGMV. Multiple comparisons were corrected for across regional volumes (i.e., 68 comparisons).

## Acknowledgement

Data used in the preparation of this article were obtained from the Adolescent Brain Cognitive DevelopmentSM (ABCD) Study (https://abcdstudy.org), held in the NIMH Data Archive (NDA). This is a multisite, longitudinal study designed to recruit more than 10,000 children age 9-10 and follow them over 10 years into early adulthood. The ABCD Study® is supported by the National Institutes of Health and additional federal partners under award numbers U01DA041048, U01DA050989, U01DA051016, U01DA041022, U01DA051018, U01DA051037, U01DA050987, U01DA041174, U01DA041106, U01DA041117, U01DA041028, U01DA041134, U01DA050988, U01DA051039, U01DA041156, U01DA041025, U01DA041120, U01DA051038, U01DA041148, U01DA041093, U01DA041089, U24DA041123, U24DA041147. A full list of supporters is available at https://abcdstudy.org/federal-partners.html. A listing of participating sites and a complete listing of the study investigators can be found at https://abcdstudy.org/consortium_members/. ABCD consortium investigators designed and implemented the study and/or provided data but did not necessarily participate in the analysis or writing of this report. This manuscript reflects the views of the authors and may not reflect the opinions or views of the NIH or ABCD consortium investigators.

Data used in the preparation of this article were obtained from the Adolescent Brain Cognitive Development (ABCD) Study (https://abcdstudy.org), held in the NIMH Data Archive (NDA). The ABCD Study is supported by the National Institutes of Health and additional federal partners. This manuscript reflects the views of the authors and may not reflect the opinions or views of the NIH or ABCD consortium investigators. The preparation of this manuscript was also partially supported by funding from the National Institute of Mental Health to Waller (R01MH125904) and Alexander-Bloch, Seidlitz, and Dorfschmidt (R01MH133843).

