## Supplementary_Materials for "Comparing Brain-Behavior Relationships Across Dimensional, Tail-Sampled, and Propensity-Matched Models"

**Figure S1.** Standardized mean differences (SMD) for unadjusted vs. adjusted samples

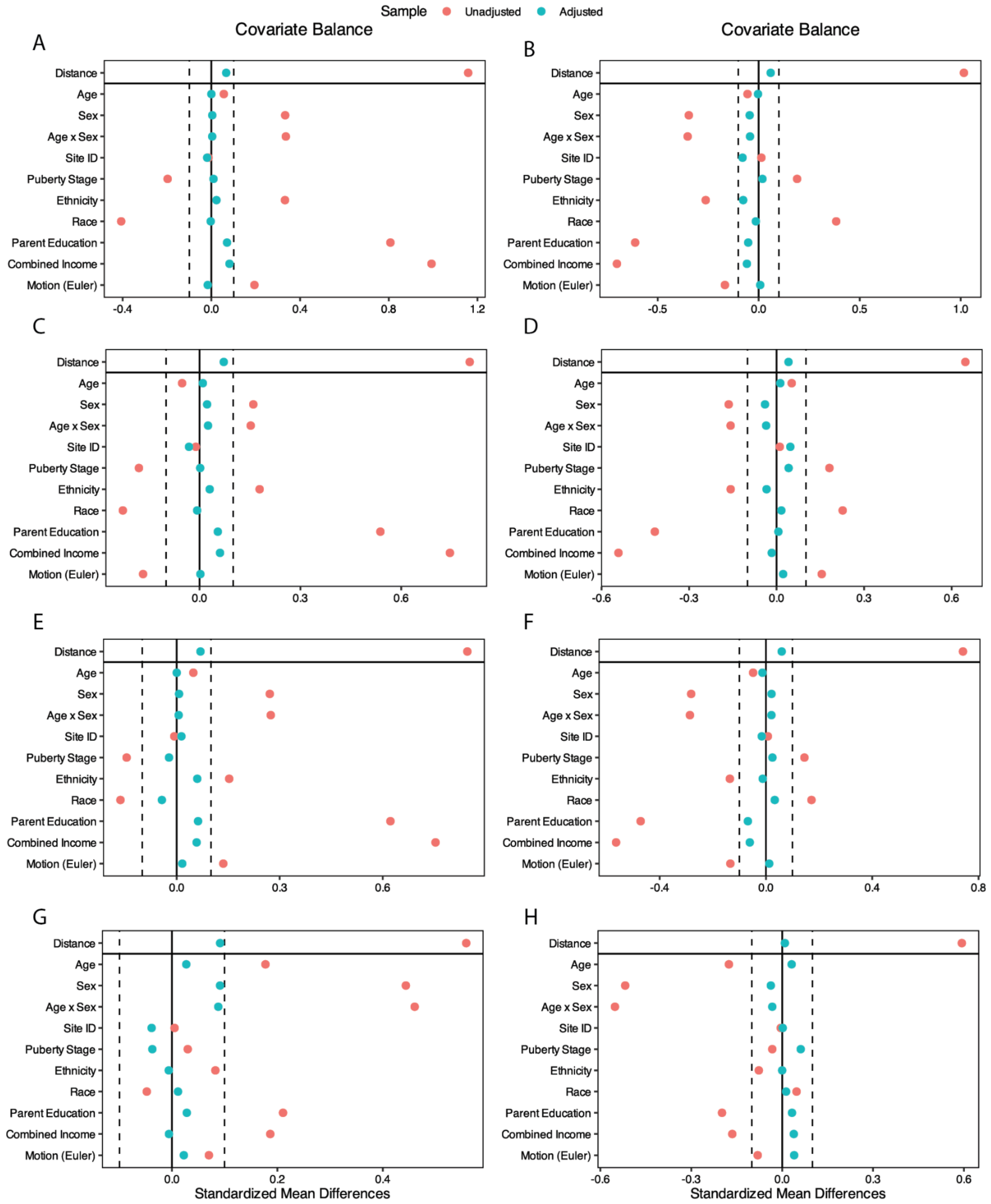

**Note.** Standardized mean differences (SMD) for unadjusted vs. adjusted samples were reduced with propensity matching. (**A**) SMD for treatment = high cGM, and (**B**) low cGM. (**C**) SMD for treatment = high WM, and (**D**) low WM. (**E**) SMD for treatment = high sGM, and (**F**) low sGM. (**G**) SMD for treatment = high CSF, and (**H**) low CSF.

**Figure S2.** Robust Effect Size Index (RESI) between cGM and example biological, cognitive, and clinical outcomes

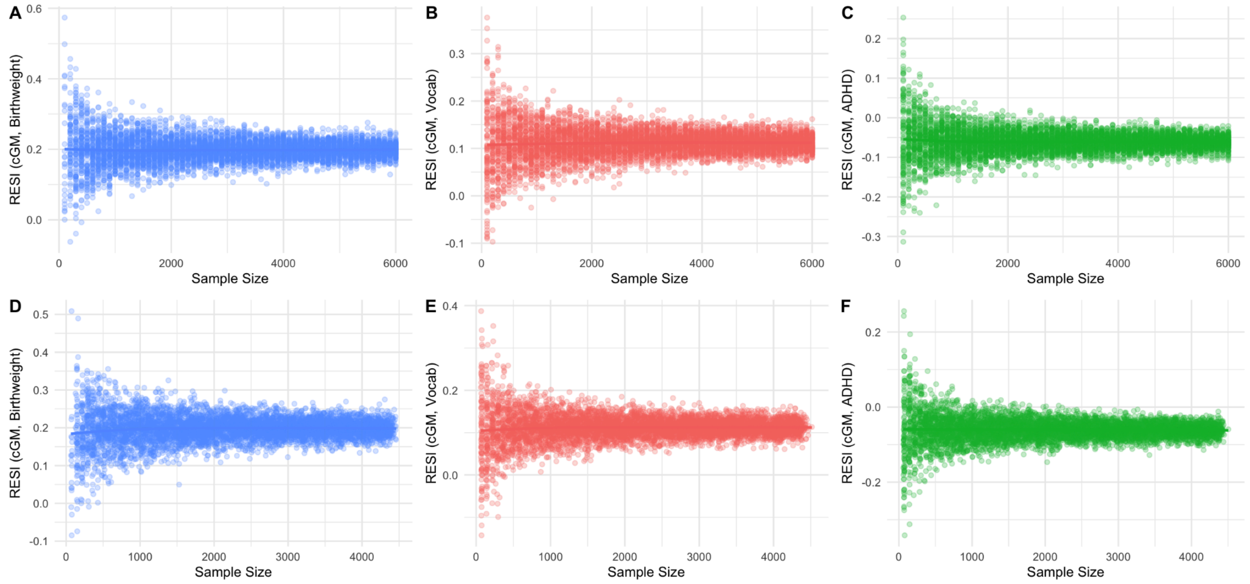

**Note**. Robust Effect Size Index (RESI) between cGM and example biological, cognitive, and clinical outcomes exhibited overall stability as sample size increased. In dimensional models (A-C) and in median absolute deviation models (D-F), RESI variability decreased and estimate stabilized, converging to a consistent central value.

**Figure S3.** RESI values representing the relationship between regional cGM and biological outcomes.

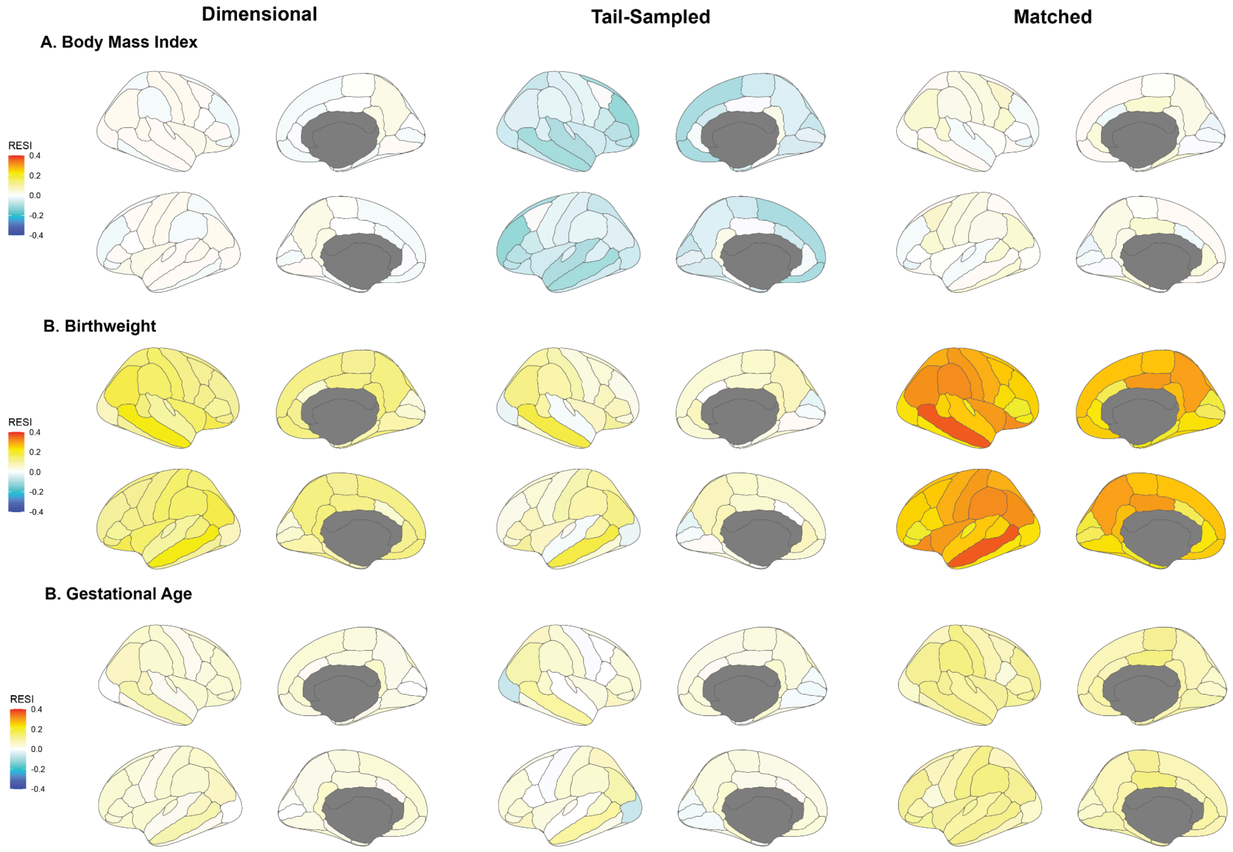

**Note.** RESI values representing the relationship between regional cGM and (**A**) body mass index, (**B**) birthweight, and (**C**) gestational age, across dimensional (**left**), tail-sampled (**center**), and propensity matched (**right**) models.

**Figure S4.** RESI values representing the relationship between regional cGM and cognitive outcomes.

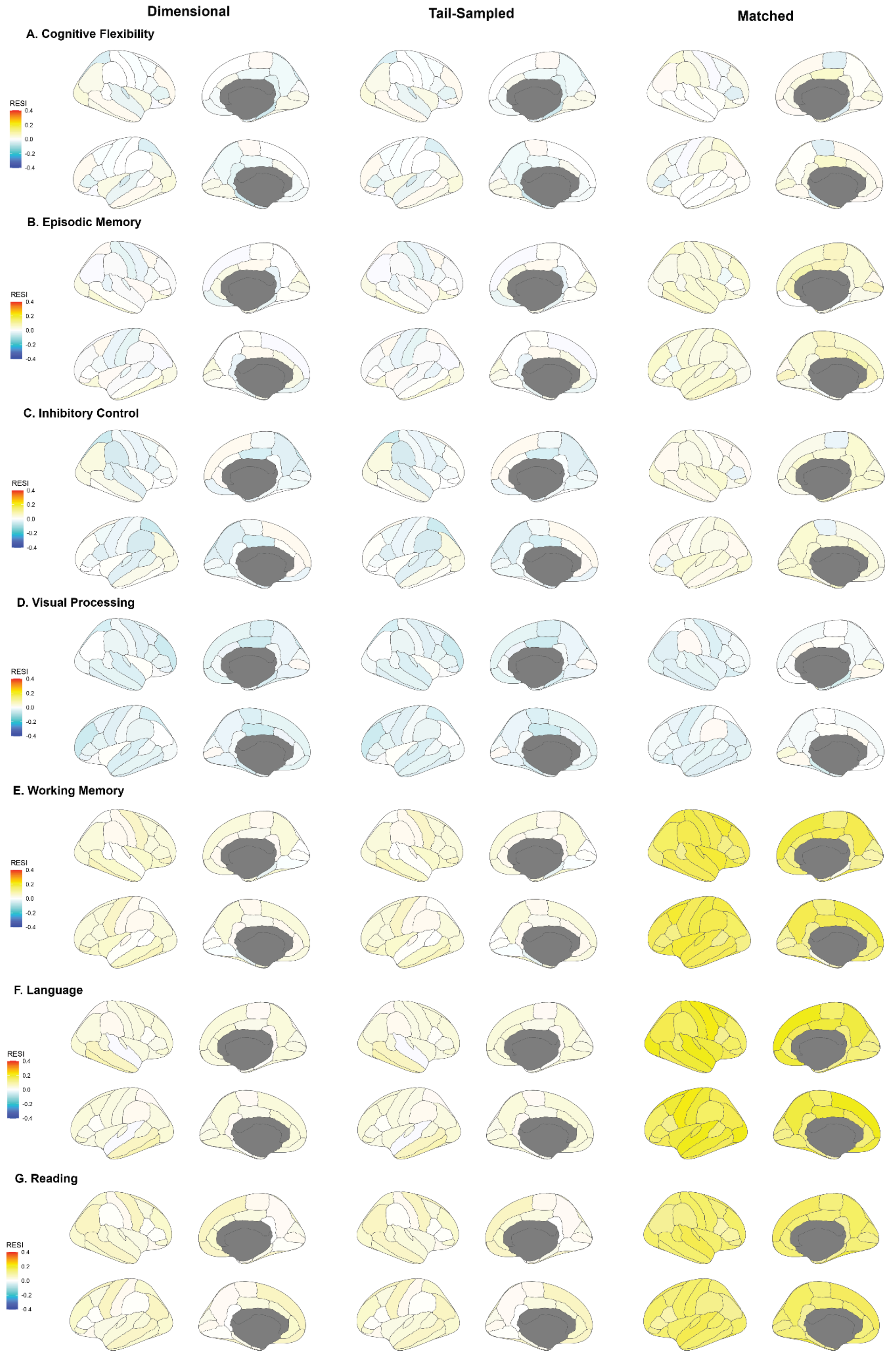

**Note.** RESI values representing the relationship between regional cGM and (**A**) cognitive flexibility, (**B**) episodic memory, (**C**) inhibitory control, (**D**) visual processing, (**E**) working memory, (**F**) Language, and (**G**) Reading across dimensional (**left**), tail-sampled (**center**), and propensity matched (**right**) models.

**Figure S5.** RESI values representing the relationship between regional cGM and clinical outcomes.

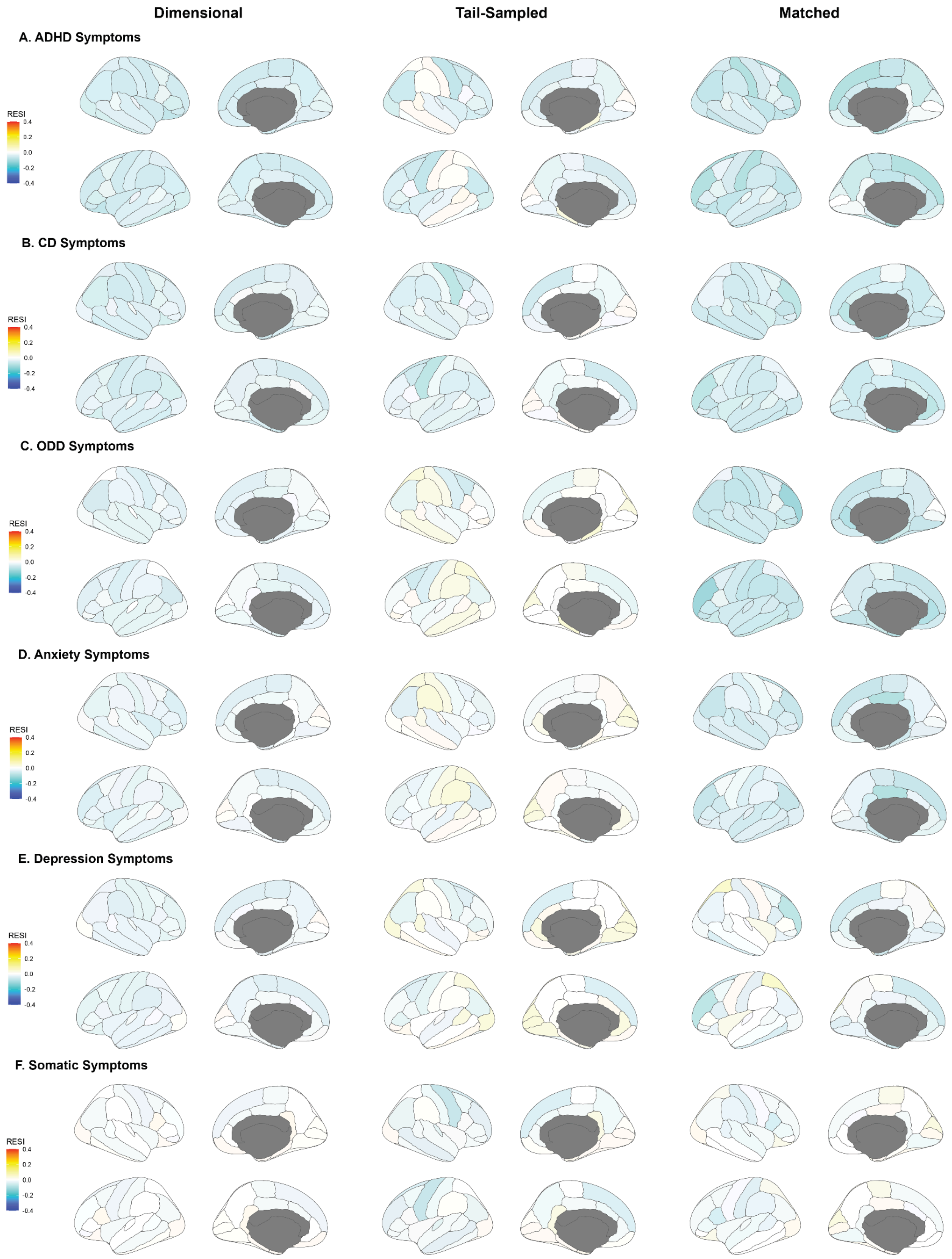

**Note**. RESI values representing the relationship between regional cGM and (**A**) ADHD Symptoms, (**B**) CD Symptoms, (**C**) ODD Symptoms, (**D**) Anxiety Symptoms, (**E**) Depression Symptoms, and (**F**) Somatic Symptoms across dimensional (**left**), tail-sampled (**center**), and propensity matched (**right**) models.

**Figure S6.** Sample selection flow chart for each tissue type

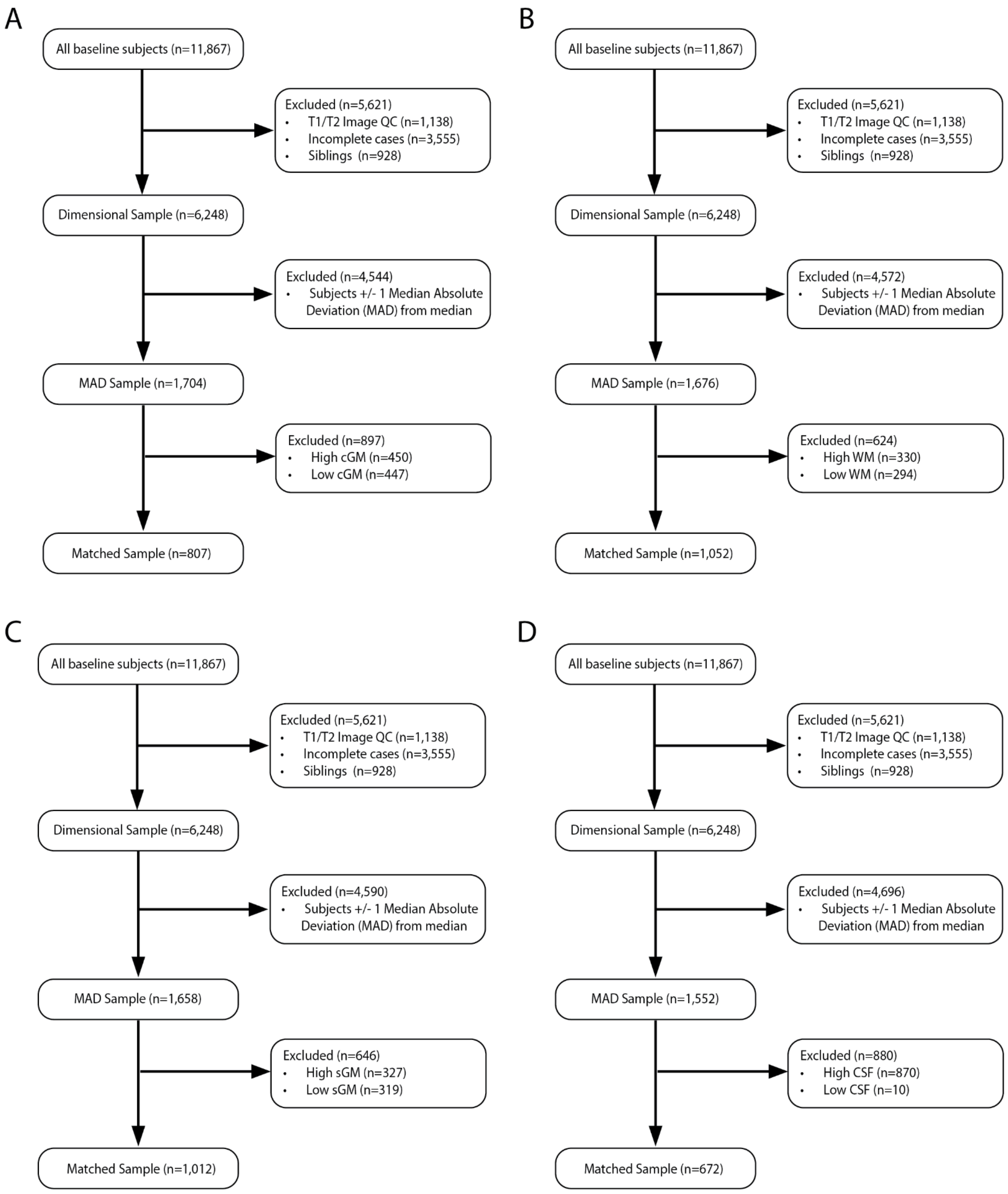

**Note.** Sample selection flow chart for each tissue type, (**A**) cGM, (**B**) WM, (**C**) sGM, (**D**) CSF.

**Figure S7.** Differences between complete and incomplete data.

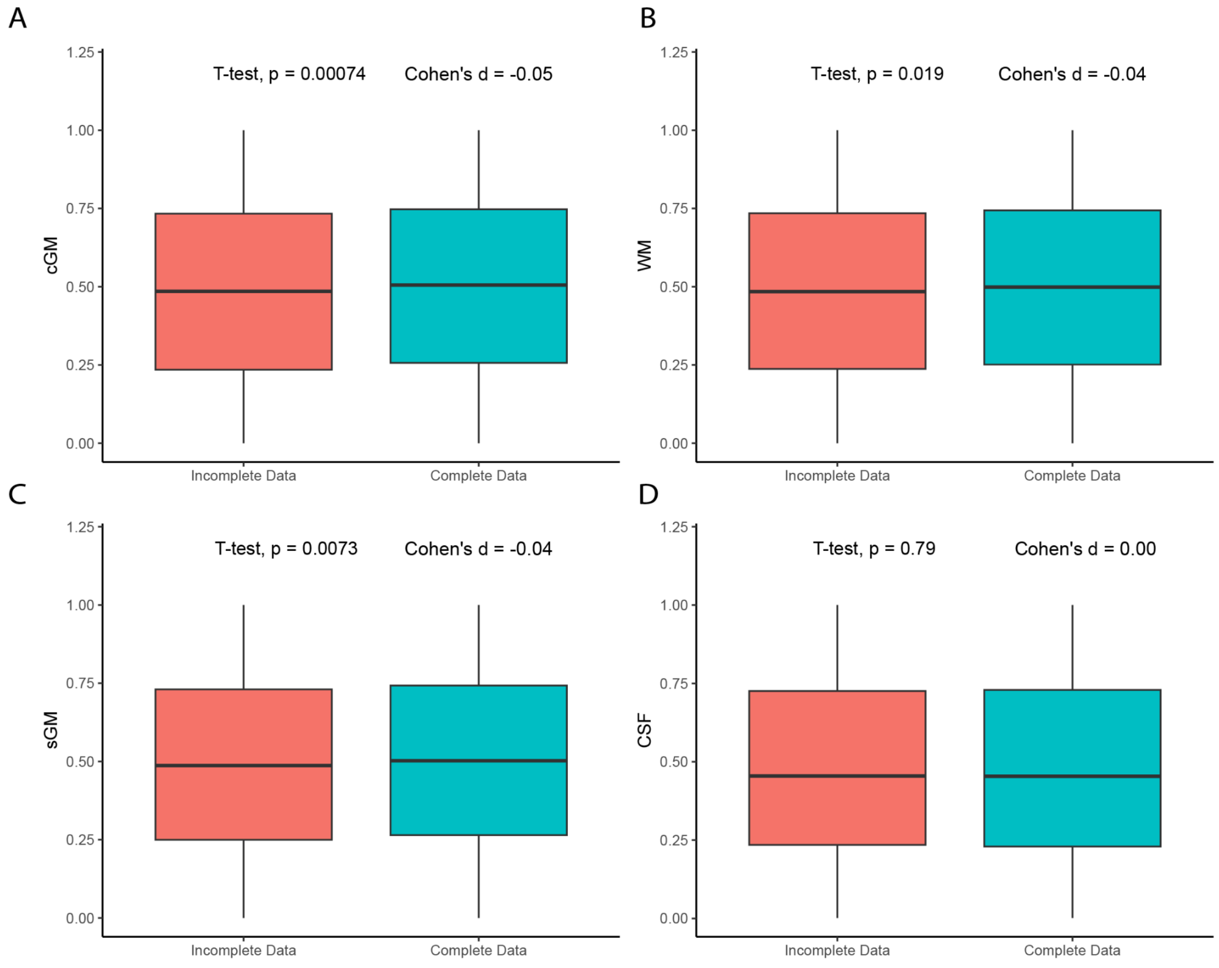

**Note.** Differences between complete and incomplete data. (**A**) Participants with missing data differed significantly on their average centile score for cGM from participants with complete data, but effect sizes were negligible. (**B**) Participants with missing data differed significantly on their average centile score for WM from participants with complete data, but effect sizes were negligible. (**C**) Participants with missing data differed significantly on their average centile score for sGM from participants with complete data, but effect sizes were negligible. (**D**) Participants with missing data did not differ significantly on their average centile score for CSF from participants with complete data.

**Appendix S1.** Comparison of Matched, Tail-sampled, and Dimensional Samples on matching covariates

Groups matched on cGMV did not differ significantly on distributions of sex, ethnicity, or measures of age, pubertal stage, or motion (**Table S1**). All groups differed significantly on household income, with tail-sampled high cGMV groups having the highest values and tail-sampled low groups having the lowest values. Patterns of results were similar for parental education. Interestingly, matched high groups both differed significantly from the average group on both measures, where matched low groups had higher income and parental education, and matched high groups had lower income and parental education.

Groups matched on sGMV did not differ significantly on distributions of ethnicity or measures of age, pubertal stage, parental education, or motion (**Table S2**). Groups differed significantly on measures of parental education and household income, where tail sampled high sGMV groups had the highest levels and tail-sampled low had the lowest levels. Once again, matched low groups had higher income and parental education, and matched high groups had lower income and parental education, though matched groups did not differ significantly from the average group.

Groups matched on WMV did not differ significantly on distributions of sex or ethnicity, or measures of age, pubertal stage, parent education, household income, or motion (**Table S3**). Groups differed significantly on measures of parental education and household income, where tail sampled high WMV groups had the highest levels and tail-sampled low had the lowest levels. Once again, matched low groups had higher income and parental education, and matched high groups had lower income and parental education.

Finally, groups matched on CSF did not differ significantly on distributions of sex, ethnicity, and race, or measures of age, pubertal stage, parent education, or motion (**Table S4**). Groups differed significantly on measures of parental education and household income, where tail sampled high CSF groups had the highest levels and tail-sampled low had the lowest levels. Once again, matched low groups had higher income and parental education, and matched high groups had lower income and parental education, though matched groups did not differ significantly from the average group.

**Table S1.** Demographic information for samples derived for cGM models and ANOVA comparisons.
**Table S2.** Demographic information for samples derived for sGM models and ANOVA comparisons.
**Table S3.** Demographic information for samples derived for WM models and ANOVA comparisons.
**Table S4.** Demographic information for samples derived for WM models and ANOVA comparisons.
**Table S5.** Regression results for models predicting biological outcomes using cGM across 3 modeling frameworks, controlling for covariates
**Table S6.** Regression results for models predicting biological outcomes using sGM across 3 modeling frameworks, controlling for covariates
**Table S7.** Regression results for models predicting biological outcomes using WM across 3 modeling frameworks, controlling for covariates
**Table S8.** Regression results for models predicting biological outcomes using CSF across 3 modeling frameworks, controlling for covariates
**Table S9.** Regression results for models predicting biological outcomes using regional cGM across 3 modeling frameworks, controlling for covariates
**Table S10.** Regression results for models predicting cognitive outcomes using cGM across 3 modeling frameworks, controlling for covariates
**Table S11.** Regression results for models predicting cognitive outcomes using sGM across 3 modeling frameworks, controlling for covariates
**Table S12.** Regression results for models predicting cognitive outcomes using WM across 3 modeling frameworks, controlling for covariates
**Table S13.** Regression results for models predicting cognitive outcomes using CSF across 3 modeling frameworks, controlling for covariates
**Table S14.** Regression results for models predicting cognitive outcomes using regional cGM across 3 modeling frameworks, controlling for covariates
**Table S15.** Regression results for models predicting clinical outcomes using cGM across 3 modeling frameworks, controlling for covariates
**Table S16.** Regression results for models predicting clinical outcomes using cGM across 3 modeling frameworks, controlling for covariates
**Table S17.** Regression results for models predicting clinical outcomes using WM across 3 modeling frameworks, controlling for covariates
**Table S18.** Regression results for models predicting clinical outcomes using CSF across 3 modeling frameworks, controlling for covariates
**Table S19.** Regression results for models predicting clinical outcomes using regional cGM across 3 modeling frameworks, controlling for covariates
